# Integration of proteogenomic analyses in esophageal squamous cell carcinoma

**DOI:** 10.64898/2026.04.20.719529

**Authors:** Guixue Hou, Shaohang Xu, Fang Zhao, Lijuan Duan, Haijun Yang, Junkuo Li, Ying Hu, Siqi Liu

**Author notes:** Corresponding authors: **Siqi Liu**, BGI-SHENZHEN, Shenzhen, Guangdong, China, 518083, **Fuyou Zhou**, Anyang Key Laboratory for Esophageal Cancer Research, Anyang Cancer Hospital, the Forth Affiliated Hospital of Henan University of Science and Technology, Anyang, Henan, China, 455001, **Ying Hu**, School of Life Science and Technology, Harbin Institute of Technology, Harbin, Heilongjiang Province 150001, China.

## Abstract

Esophageal squamous cell carcinoma (ESCC) is still lack of clinically molecular subtyping and effective therapeutic strategies. Herein, a total of 46 paired tissue samples of esophageal squamous cell carcinoma (ESCC) were collected and subjected to a systematic proteogenomic evaluation. Consensus assessment of the ESCC-related transcriptomes and TCGA dataset revealed several consensual modes of gene expression related to ESCC specificity, with 8 plasma-detectable hub proteins that could discriminate ESCC from others. Three ESCC molecular subtypes were defined and validated based on proteome data, including pCC1 with activated immune response and best survival outcome, pCC2 as cell cycle subtype with relative worse outcome, and pCC3 with worst outcome that expressed more cell adhesion related proteins. Furthermore, we proposed potential therapeutic strategies for improving survival outcomes in patients with different ESCC molecular subtypes. This integrative proteogenomic analysis provided a novel view of ESCC-dependent molecular information.

## Introduction

Genomic analysis of esophageal squamous cell carcinoma (ESCC) has led to substantial progress in the characterization of the genomic landscape of this cancer. The following key findings have been achieved: 1) significantly mutated genes, such as TP53, FAT1, NOTCH1, PIK3CA, PTEN and DCDC1^1–4^; 2) different frequent gene amplifications (CCND1, SOX2, TP63, ERBB2, VEGFA, GATA4 and GATA6) between esophageal adenocarcinomas and squamous cell carcinomas^5^; and 3) the dysregulation of multiple pathways, such as RTK-MAPK-PI3K, the cell cycle and epigenetic regulation^6,7^. Determining how genetic aberrations related to ESCC drive tumor phenotypes at the molecular level is critical in the study of oncogenic mechanisms.

Proteogenomic analysis is an emerging approach that combines genomics and proteomics at the multi-omics level using high-throughput sequencing and mass spectrometry technology. This approach is promising for advancing translational and clinical research. Jin et al conducted a comprehensive proteogenomic study involving the transcriptomic, proteomic and phosphoproteomic characterization of 24 paired ESCC and corresponding adjacent tissue samples. This integrative analysis revealed that ESCC tissues had distinct signatures at the transcriptome, proteome and phosphoproteome levels versus normal tissues, whereas there was a striking discordance between mRNAs and proteins at a median correlation value of 0.07, implying different regulation at the two expression levels in ESCC^8^. Recognizing the volume of data reported in the literature and the lack of a comprehensive summary, Tungekar et al. compiled the proteogenomic information available about ESCC and developed a new database, ‘ESCC ATLAS’, that catalogs all the genes and related molecular signatures involved in ESCC based on an exhaustive literature survey^9^. Recently, the ESCC-related posttranslational modifications (PTM) study was carried out using a multi-omics approach, including an analysis of the phosphorylome, lysine acetylome and succinylome strongly associated with ESCC^10,11^. While proteogenomic analysis has provided unique insights into the molecular mechanisms underlying ESCC, several key issues remain unresolved. Firstly, integrative strategies to assess the global responses of ESCC-related genes should be explored, not only restricted within analysis of differentially expressed genes (DEGs) or differentially expressed proteins (DEPs). Although the expression abundance of RNA and proteins is generally discordant, integration of the two datasets is required for molecular signatures related with ESCC. Secondly, the molecular specificity and mechanism of ESCC is still not clear, and the integration of ESCC with public cancer datasets is essential. Thirdly, the study about ESCC molecular subtypes and therapeutic recommendation is limited, and it is important to acquire and accumulate more datasets especially at multi-omics level.

To address the issues raised above, we conducted a proteogenomic study of ESCC, in which 46 paired ESCC surgery samples were collected and an RNA-seq and LC□MS/MS based proteogenomic approach was used to globally and quantitatively investigate ESCC-related gene expression. We tried to integrate these datasets in several aspects, 1) several consensual modes of gene expression related to ESCC specific, pan-gastrointestinal or pan-squamous cancer were defined by consensus assessment with ESCC-related transcriptomes, proteomes, TCGA^12^ and CPTAC^13,14^ datasets. 2) ESCC-specific module with 8 plasma-detectable hub proteins that could discriminate ESCC from other types of cancer were categorized; 3) three ESCC molecular subtypes were classified based on proteome data, which was beneficial for survival outcome prediction and therapeutic guidance for ESCC patients. Overall, this integrative proteogenomic analysis provided a novel view of ESCC-dependent molecular information.

## Results

### The proteogenomic atlas of ESCC paired tissues

ESCC and corresponding adjacent tissues were collected from 46 ESCC patients (the patients’ clinical information is summarized in **Supplementary Data 1**). mRNA and proteome data were acquired from these tissues via RNA-seq and LC□MS/MS in data-independent acquisition (DIA) mode (**Fig.1a**). At the transcriptome level, RNA-seq identified 18,799 genes across 40 paired ESCC samples (**Supplementary Data 2**), with an average of 17,130 genes/sample (detailed mRNA numbers for each sample are shown in **Supplementary Fig.1a**). In the 46 paired ESCC tissues, 105,499 peptides corresponding to 10,066 proteins (**Supplementary Data 3**) were identified with DIA mode (detailed identified protein number for each sample depicted in **Supplementary Fig.1a**); 7,985 proteins were present in at least 20% of the samples (i.e., quantifiable proteins).

**Figure 1.**
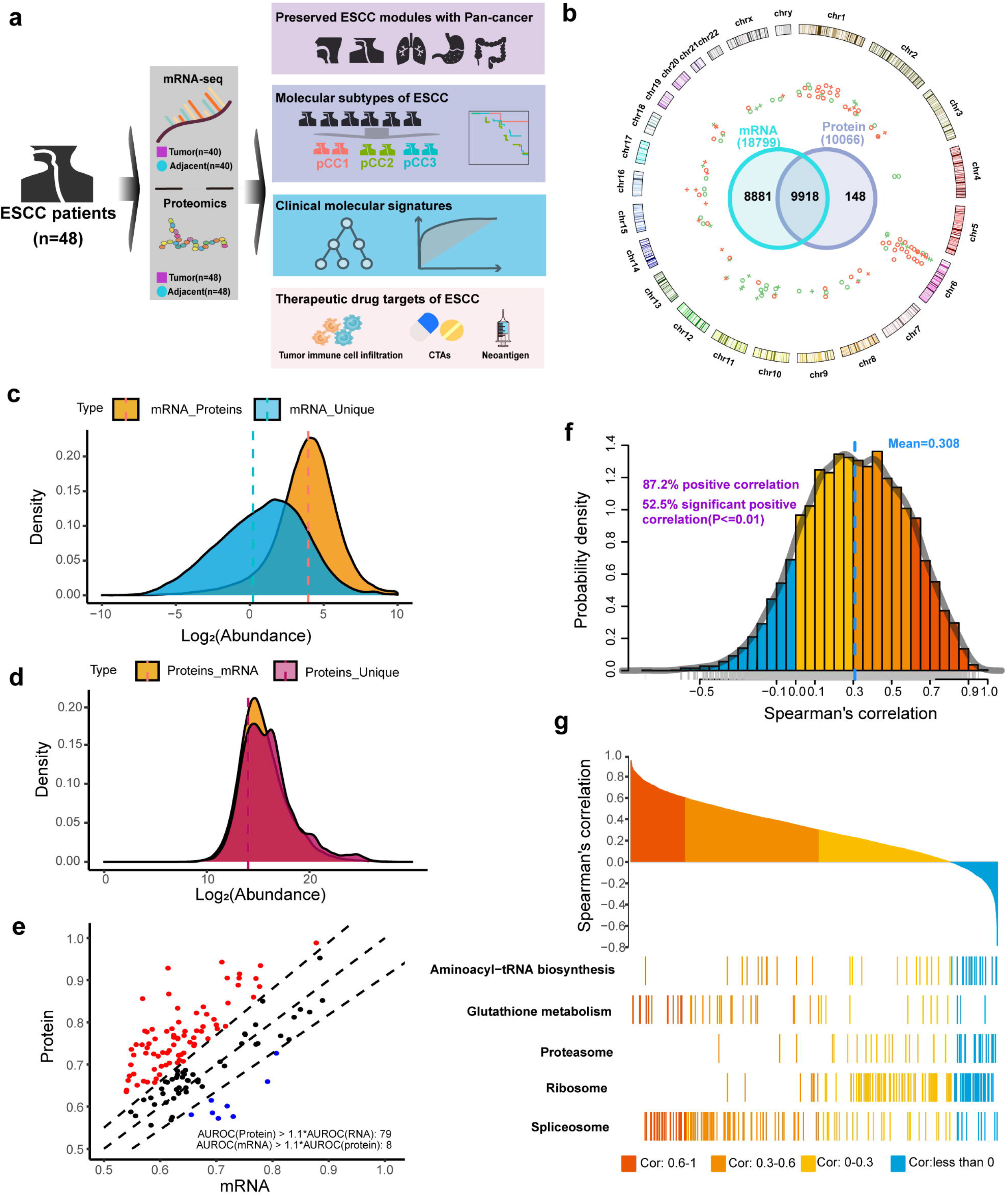
Overview of the mRNA and protein data atlas. (**a**) Study design of the proteogenomic analysis for ESCC. (**b**) Genes identified at the mRNA and protein levels. Genes only identified at the protein level were located at the chromosome locus circle with differential annotation comparing ESCC with adjacent samples. Red circles represent significantly upregulated genes in ESCC, and green circles represent significantly downregulated genes in ESCC, with stars representing nonsignificant dysregulated genes. (**c**) Abundance distribution for genes co-identified at both the mRNA and protein levels and genes identified only at the mRNA level. (**d**) Abundance distribution for genes co-identified at both the mRNA and protein levels and genes identified only at the protein level. (**e**) Density distribution of Spearman’s correlation between mRNA and proteins. The mean transcript-protein pair correlation, 0.3, is consistent with other proteogenomic studies. (**f**) Area under the receiver operating characteristic curve (AUROC) for KEGG (Kyoto Encyclopedia of Genes and Genomes) pathway membership prediction using RNA and protein data. Red and blue indicate pathways with >10% difference between the two. (**g**) Waterfall plot of representative pathways related to different correlation coefficients for transcript-protein pairs.

There were 9,918 genes co-identified in the transcriptome and proteome datasets, while 8,881 genes were only detected by RNA-seq and 148 genes were uniquely perceived by LC-MS/MS. For these genes whose protein products were only identified by LC-MS/MS, their chromosomal locations are presented in **Fig.1b**; many of these genes are enriched in the MHC regions of chromosome 6 (20/148), and some were located on chromosomes 3, 4 and 5, suggesting that MHC-based immune responses are involved in ESCC. The mRNA levels for these genes whose expressed products were only found by RNA-seq or co-identified by RNA-seq and LC□MS/MS are shown in **Fig.1c**. The mRNA levels of the co-identified genes were higher than those identified only by RNA-seq, whereas the protein level of the co-identified genes versus those uniquely detected by LC□MS/MS were similar as illustrated in **Fig.1d**, revealing that the genes identified only by proteomics were expressed at general level. The OmicsEV^15^ analyses of the ESCC-proteogenomic data are illustrated in **Fig.1e**. These analyses revealed that the distribution patterns of the area under the receiver operating characteristic curve (AUROC) obtained from the protein data were substantially different from those obtained from the mRNA data. Seventy-nine function terms were predicted by proteins with outperformance; only 8 terms were predicted from the mRNA dataset. Thus, protein data are likely better for annotating the functions of ESCC-related genes.

Principal component analysis clearly separated the ESCC tumor samples and corresponding adjacent tissues based on either mRNA or protein data (**Supplementary Fig.1b and 1c**). Differential analyses of the transcriptome and proteome revealed that there were 5,102 upregulated and 537 downregulated DEGs, and 2,326 upregulated and 116 downregulated DEPs (**Supplementary Fig.1d**) in the ESCC tissues compared with adjacent ones. However, the gene-wise correlation between proteomics and RNA-Seq data was poor (mean correlation coefficient = 0.308, **Fig.1f**). Using Spearman’s analysis towards the abundance correlations, the co-identified genes were broadly categorized to four functional groups (**Fig.1g**); those with high correlation were enriched in glutathione metabolism and spliceosome functions, whereas those with diverse correlation were annotated to aminoacyl-tRNA biosynthesis, proteasome and ribosome functions. Obviously, the ESCC-dependent characteristics of mRNAs and proteins are distinct and unlikely to be substitutable with each other.

### WGCNA of the transcriptomes and proteomes of ESCC paired tissues

With strict criteria for the genes co-identified by RNA-seq and MS/MS in more than 50% of esophageal tissues, 5,480 genes were used for WGCNA^16,17^ to further investigate the abundance trends between ESCC and adjacent tissues. By setting a small minimum WGCNA module size of 30 genes/proteins with a low propensity to merge modules (merge height of 0.07), a total of 26 modules (**Supplementary Data 4**) were established: ME1 (largest, 789 genes/proteins) to ME26 (smallest, 37 genes/proteins). As illustrated in **Supplementary Fig.2**, the genes in most modules were significantly enriched in specific biological processes and cellular functions.

The module-trait relationships containing the correlations between the transcriptome or proteome in ESCC and adjacent tissues and the correlation consensus based on these WGCNA modules are shown in **Fig.2a**. Among these modules, 14 (ME1, ME2, ME3, ME4, ME7, ME11, ME15, ME18, ME19, ME22, ME23, ME25 and ME26) demonstrated upregulation at both the mRNA and protein levels in ESCC. Only ME5 was downregulated at both expression levels in ESCC. These 15 modules are termed shared modules. There were 7 modules (ME6, ME12, ME13, ME14, ME16, ME24 and ME20) with upregulated expression and one module (ME21) with downregulated expression at the mRNA level but not at the protein level, while there were 4 modules (ME8, ME9, ME10 and ME17) with downregulated expression at the protein level but not at the mRNA level. These 11 modules are referred to as mRNA- or protein-specific modules. Pearson correlation was further implemented to assess the abundance correlation of different expression levels in these modules. The Pearson coefficients of the shared modules reached 0.69 (**Fig.2b**), indicating a moderate correlation of mRNA and protein abundance changes in these modules; in the mRNA- or protein-specific modules, the Pearson coefficients were −0.06, indicating no correlation between mRNA and protein levels in these modules in ESCC (**Fig.2c**). GSEA^18,19^ was used to explore the gene set enrichment of hallmarks in these ESCC-related modules. As shown in **Fig.2d**, several representative hallmarks were specifically enriched in the shared modules, mTORC1 signaling and E2F targets in ME1, E2F and MYC targets in ME2, the G2/M checkpoint in ME3, and epithelial-mesenchymal transition in ME7 and ME15. Hence, these representative hallmarks are likely to be ESCC biomarkers that can be measured at either the transcriptional or translational level.

**Figure 2.**
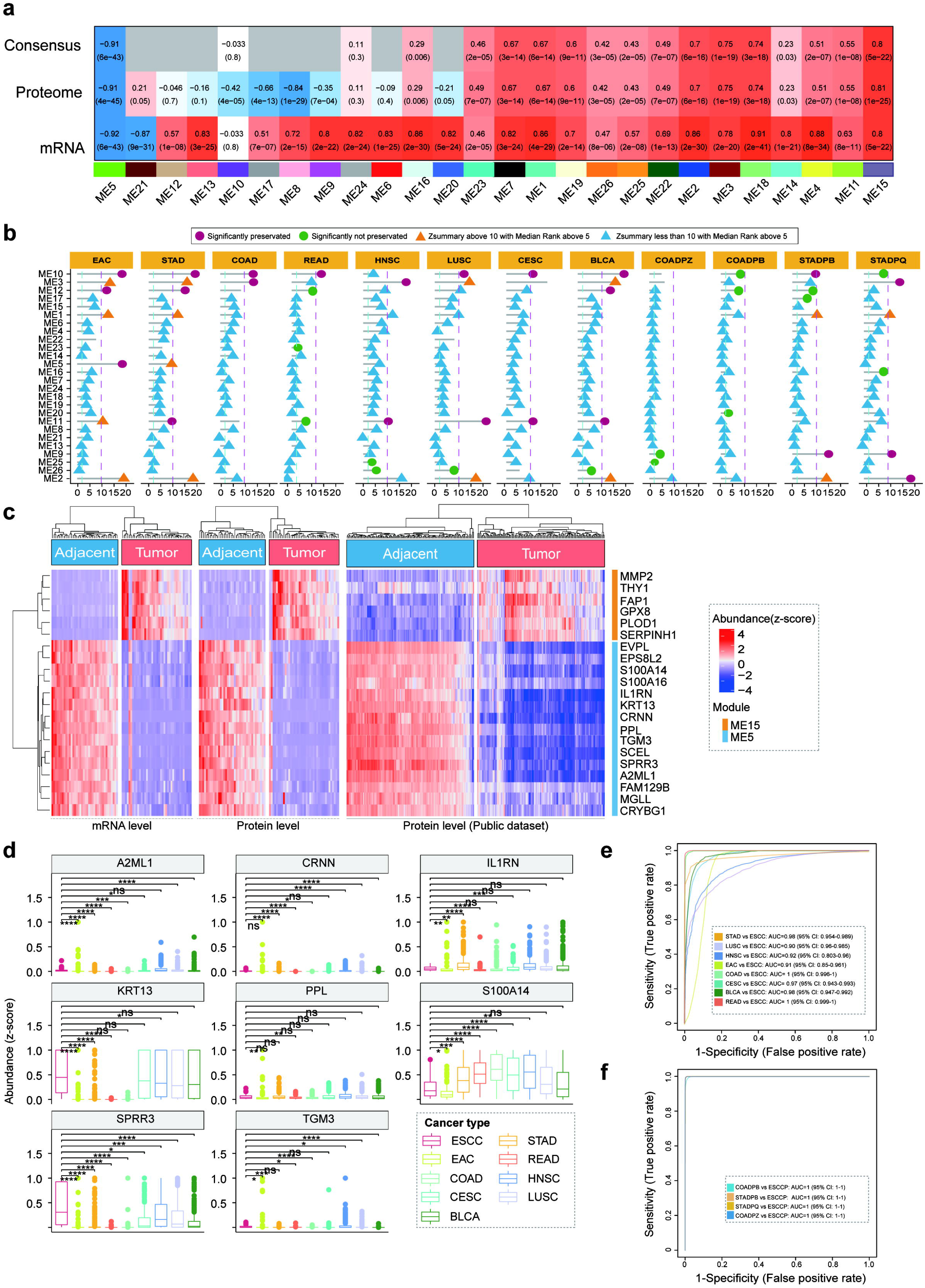
WGCNA using mRNA and proteome data for ESCC. (**a**) Consensus WGCNA using ESCC mRNA and proteome data. Each col corresponds to a module eigengene, each column to a trait. Each cell contains the corresponding correlation and p value. The table was color-coded by the correlation between modules and ESCC. (**b**) Correlation of fold changes for genes quantified in consisted modules between mRNA and proteins. (**c**) Correlation of fold changes for genes quantified in inconsistent modules between mRNA and proteins. (**d**) Heatmap of genes in modules with representative hallmarks at the transcriptome and proteome levels.

### ESCC signatures derived from ESCC-related modules and hub genes

Laird et al. used TCGA data and found that esophageal carcinomas could be broadly grouped into two categories, pan-gastrointestinal (Pan-GI) cancers, including the subcategories of esophageal adenocarcinoma (EAC), stomach adenocarcinoma (STAD), colon adenocarcinoma (COAD) and rectum adenocarcinoma (READ), and Pan-Squamous cancers, including the subcategories of head and neck squamous cell carcinoma (HNSC), lung squamous cell carcinoma (LUSC), cervical squamous cell carcinoma and endocervical adenocarcinoma (CESC) and bladder urothelial carcinoma (BLCA)^20^. A preservation assessment of the pan-carcinoma features based on gene expression can be judged by two critical parameters, Zsummary and MedianRank^21^; a strong preservation of modules is assumed with a Zsummary value greater than 10 and a MedianRank value less than 5. The transcriptome data from the ESCC-related modules were integrated into an assessment with Zsummary and MedianRank (**Fig.3a**). As shown, the transcription status of the ME10 genes was preserved in all Pan-GI and Pan-Squamous cancers except for HNSC, while that of the ME11 genes was preserved in the Pan-Squamous HNSC, LUSC and CESC and the Pan-GIs STAD and BLCA. In addition, ME3 transcription was preserved in COAD, HNSC and CESC. A total of 12 modules were highly preserved in ESCC, including ME4, ME5, ME6, ME7, ME8, ME13, ME14, ME15, ME18, ME19, ME21 and ME24. Importantly, the transcription status of ME5 was specifically preserved in ESCC and EAC, while that of ME15 was only ESCC-specific.

**Figure 3.**
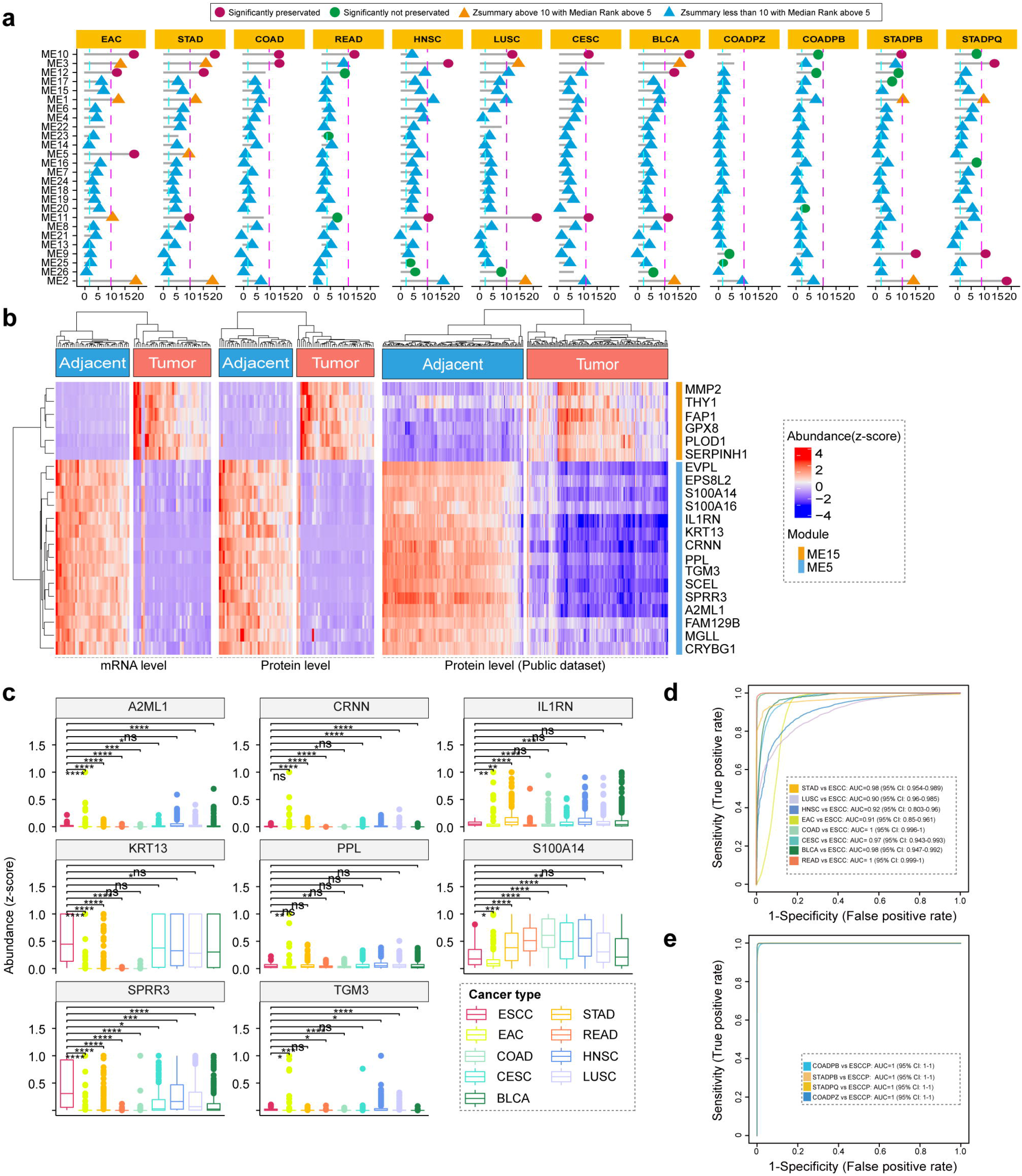
Pan-cancer analysis for preserved modules related to ESCC. (**a**) Preservation of ESCC network modules in pan-cancer datasets. (**b**) Heatmap of the mRNA abundance and protein abundance distribution for hub genes in ESCC-specific modules ME5 and ME15 among ESCC and adjacent samples. (**c**) Boxplot of the protein abundance distribution for 8 hub genes in different cancer types. Receiver operating characteristic (ROC) curves for the essential gene prediction classifiers on ESCC with other cancer types in mRNA (**d**) or protein datasets (**e**).

For preservation assessment at proteome level, four available published cross-carcinoma proteome datasets including gastric and colon cancers were selected and represented as COADPZ^22^, COADPB^23^, STADPQ^24^ and STADPB^25^. The preservation analysis between ESCC and gastric or colon cancers at the proteome level revealed no preserved modules between ESCC and colon cancers; however, three modules, ME2, ME3 and ME9, were preserved between ESCC and gastric cancer (**right four columns on Fig.3a**). Moreover, integration of Zsummary and MedianRank for the proteomes in the ESCC-related modules suggested that ME5 and ME15 are highly ESCC-dependent modules. Intriguingly, as shown in **Fig.2a**, only ME5 and ME15 demonstrate downregulated or upregulated genes with high significant correlation coefficients in ESCC (p value<0.01). The correlation characteristics mentioned above suggested that these two modules might contain ESCC-specific hub genes.

There were 294 and 141 genes in ME5 and ME15, respectively. The hub genes were estimated by setting the top 5% of genes in a module with module membership (kME) above 0.8^26^. A total of 21 hub genes were found in the two modules, including 6 in ME15 (SERPINH1, GPX8, THY1, MMP2, PLOD1 and FAP; **Fig.3b upper**) and 15 in ME5 (PPL, SPRR3, S100A14, FAM129B, CRNN, TGM3, A2ML1, EPS8L2, EVPL, SCEL, S100A16, IL1RN, CRYBG1, KRT13 and MGLL; **Figure 3b lower**). As shown in **Supplementary Fig.3a** using Cytoscape^27^, the connectivity map strongly supported the selection of these hub genes. The mRNA or protein abundance of the 21 hub genes in the two modules was hierarchically plotted against all the paired samples. As expected, the heatmaps shown in **Fig.3b left** show that the mRNA or protein abundance distribution of these hub genes in tumors were significantly distinct from that in adjacent tissues. More importantly, clustering these hub genes in a public ESCC proteome database^28^ revealed similar results, indicating that the abundance of these proteins differed in ESCC versus in adjacent tissues (**Fig.3b, right**).

Of 444 genes with elevated expression in esophagus in Human Protein Atlas^29,30^, 94 are enriched in ME5, which are termed esophagus genes in ME5 (**EGME5**) in this study. Of the 15 hub genes in ME5, 14 are present in the network of esophagus-specific genes in **Supplementary Fig.3a** (yellow nodes). Single cell type annotation in HPA revealed that 9 genes were clustered in squamous epithelial cells, indicating that these 9 genes were proven to be esophagus squamous carcinoma markers, not only esophagus tissue specificity but also squamous epithelial cell typing. Importantly, 8 of them were plasma detectable as the potential blood screening markers for ESCC. Based on the TCGA datasets described in **Fig.3a**, the transcriptional abundance of the 8 hub genes derived from EGME5 were statistically and individually appraised for ESCC specificity in comparison with other cancer types (**Fig.3c**). The abundance of transcripts and proteins of such hub genes were further assessed for their ability to discriminate ESCC from other cancers by ROC analysis using random forests algorithm. At both the transcript (**Fig.3d**) and protein (**Fig.3e**) levels, the area under the ROC curve (AUC) above 0.90 allowed the discrimination of ESCC from other cancer type. Therefore, the hub genes identified by WGCNA of ESCC proteogenomic data are valuable gene expression signatures that can characterize ESCC and discriminate it from other cancers.

### Correlation of the diagnostic and proteogenomic information of ESCC

Clinically, ESCC is divided into three tumor grades, G1 (well differentiated, low grade), G2 (moderate grade) and G3 (poorly differentiated, high grade). According to the clinical classification, the ESCC cohort in this study comprised 11 patients in G1, 20 in G2 and 8 in G3. The abundance of EGME5 at the transcript or protein level was statistically compared among the tumor and adjacent tissues based on ESCC grade, as illustrated in **Fig.4a**. The two comparisons revealed that EGME5 gene expression in the adjacent tissues was comparable in all the samples of three grades, whereas that in the tumor tissues was typically grade-dependent. The expression levels of esophageal genes in G1 were significantly higher than those in G2 and G3; expression levels were lowest in G3 samples. Thus, transcriptional or translational EGME5 signals could partially reflect ESCC grade.

**Figure 4.**
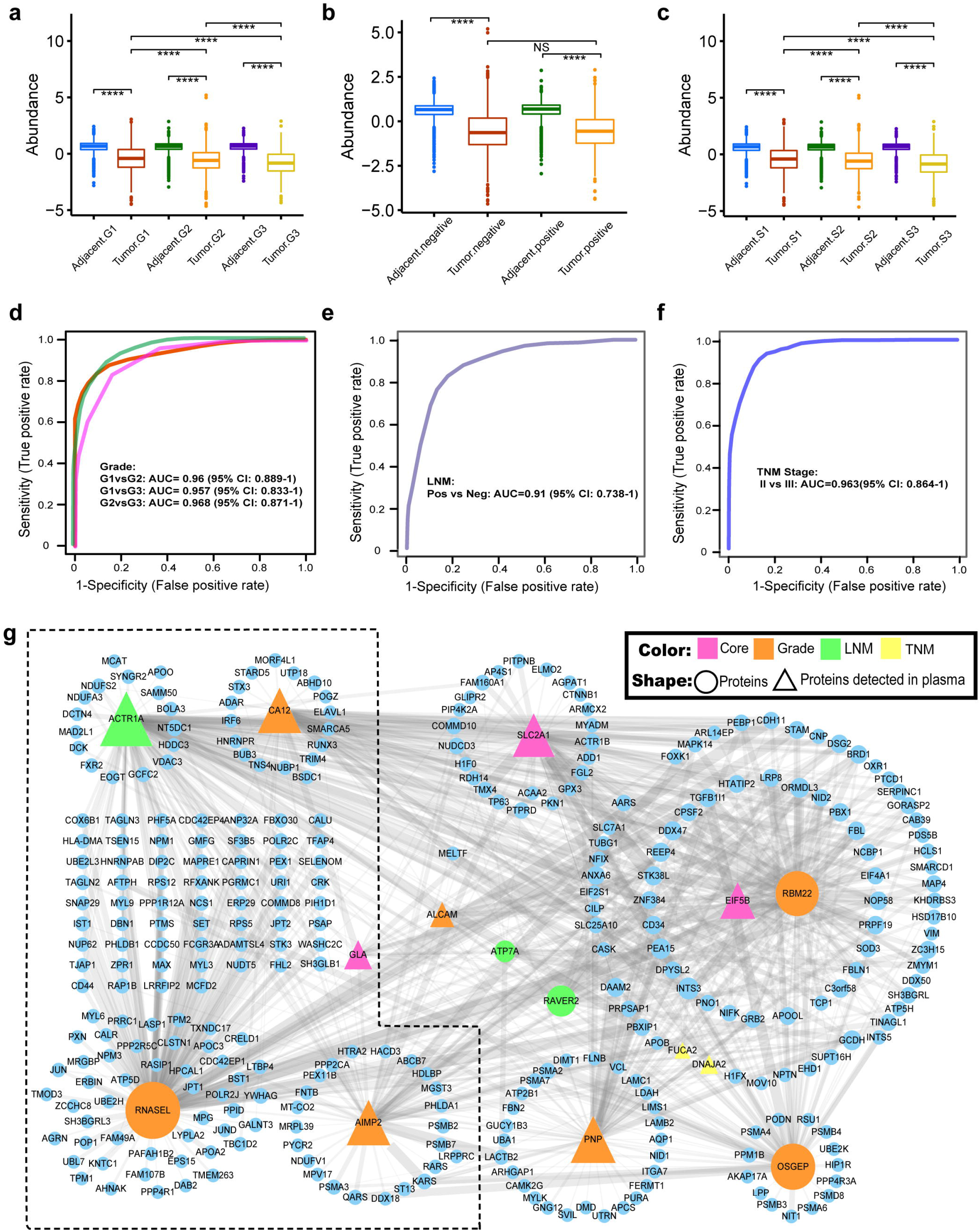
Clinical signatures related to ESCC. (**a**) Boxplot of the abundance of EGME5 in ESCC and adjacent tissues with different grades. (**b**) Boxplot of the abundance of EGME5 in ESCC and adjacent tissues from LNM-positive and LNM-negative patients. (**c**) Boxplot of the abundance of EGME5 in ESCC and adjacent tissues with TNM stages. Receiver operating characteristic (ROC) curves for the essential gene prediction classifiers on (**d**) grade, (**e**) LNM and (**f**) TNM stage datasets with combined features. (**g**) Interaction network of proteins from the biomarker signatures related to TNM stages, LNM and grades were associated and interlinked by the proteins they have in common. Protein nodes shaded in yellow were associated with different clinical pathologies. Proteins labeled with red colors represent that they could be detected in plasma by mass spectrometry.

Regarding lymph node metastasis (LNM), there were similar expression patterns of EGME5 between ESCC and adjacent groups, whereas there was no significant difference in the gene expression levels between LNM-positive and LNM-negative samples (**Fig.4b**) at either transcript or protein level. For tumor node metastasis (TNM), the abundance pattern of EGME5 expression differed among ESCC patients with TNM stage I, II and III (**Fig.4c**). However, the prediction power of EGME5 expression (transcripts or proteins) was limited, as shown in **Supplementary Fig.4a-c**. The average AUC was only 0.58 in the discrimination of ESCC with different metastasis statuses either LNM or TNM. This raised the question of whether more genes could be introduced to discriminate different ESCC metastasis statuses. The potential prediction performance using the dataset of ESCC-related DEGs or DEPs was further evaluated using the samples from ESCC patients with different tumor grades. OmicsEV^15^ was utilized to assess the degree of function prediction based on DPGs or DEPs for grade, TNM or LNM, unraveling that DEPs could achieve a better prediction for clinical parameters than DEGs (**Supplementary Fig.4 a-4j)**. Based on the DEPs and clinical information of the ESCC patients, the ROC analysis resulted in 71, 62, 21, 120 and 35 DEPs that enabled acceptable discrimination (AUC above 0.8) among tumor grades (G1 vs. G2, G1 vs. G3 and G2 vs. G3), TNM stages (II vs. III) and LNM status (positive vs. negative), respectively. Furthermore, a random forest plot was used to find the optimal combinations of these DEPs for the best discrimination. As shown in **Fig.4d**, the combination of SLC2A1, ALCAM, AIMP2, OSGEP, PNP, CA12, RBM22 and RNASEL could discriminate all grades, G1 vs. G2 and vs. G3 and G2 vs. G3, with an AUC above 0.95. As shown in **Fig.4e** and **Fig.4f**, the combination of 5 DEPs, EIF5B, GLA, ATP7A, RAVER2 and ACTR1A, could determine LNM status with an AUC of 0.91 (95% CI: 0.738-1), while the combination of 5 DEPs, FUCA2, DNAJA2, SLC2A1, EIF5B and GLA, could distinguish patients with TNM stages II and III with an AUC of 0.96 (95% CI: 0.864-1). Considering all the DEPs for predicting ESCC clinical phenotypes, some were shared with the three predictions above. For instance, EIF5B and GLA could clarify TNM stages and LNMs, and SLC2A1 could recognize TNM stages and grades, as shown in **Fig.4g**. Importantly, these DEPs could be detected in the plasma by mass spectrometry, as suggested by HPA ^29^, indicating their potential roles in diagnosing ESCC stages through plasma marker analysis.

### Molecular subtypes of ESCC defined by proteogenomic data

The DEGs and DEPs were further evaluated by consensus clustering. Based on the transcriptome data from 40 patient samples, two DEG consensus clusters (mCCs) were obtained with significant survival differences (**Supplementary Fig.5a and b**). mCC1 (18 patients) was associated with prolonged survival in comparison with mCC2 (21 patients). The DEGs in mCC1 with upregulated abundance were significantly enriched in immunity functions. Based on the proteome data from 46 patient samples, three DEP consensus clusters (pCCs) were formed. The three molecular subtypes (**Supplemental Data 5**) of ESCC identified by DEPs in **Fig.5a** show that the enriched functions in each cluster demonstrate typical protein characteristics, pCC1 (12 patients) indicating immune response, pCC2 (13 patients) reflecting the cell cycle and pCC3 (21 patients) involving focal adhesion. Comparison of the molecular subtypes of ESCC derived from DEGs and DEPs revealed that the patients in the mCC1 group were almost overlapped with those in the pCC1 group, while those in the mCC2 group were basically merged to pCC2 and pCC3. Combining the survival rates of the ESCC patients and the DEP-based molecular subtypes, we further estimated ESCC patient survival probabilities. As shown in **Fig.5b**, the subtypes of ESCC defined by DEPs were well correlated with survival outcomes, with pCC1 representing the best survival status and pCC3 indicating the worst. Evaluation of the ESCC molecular subtypes was extended to a public ESCC proteome dataset^28^ with a large cohort, and the analysis (**Fig.5c**) revealed a similar conclusion regarding survival outcomes, as indicated in **Fig.5b** that presents the three proteome-related subtypes (**Supplemental Data 5**), pCC1 (62 patients), pCC2 (33 patients) and pCC3 (29 patients). The molecular functions associated with these subtypes of ESCC are summarized in **Fig.5d**. Regarding pCC1, the upregulation of the immune system involved both MHC I and MHC II pathways^31^, JAK-STAT pathways^32^ and the CLIP complex^33^. The distribution of immune-related cell ratios estimated by Cibersoft^34^ revealed that the cells in the pCC1 subtype comprised a higher proportion of M1 macrophages (**Supplementary Fig.5d**). Regarding pCC2, cell cycle-related proteins were upregulated, including CDK1/Cyclin B, MCM and other initiation proteins. Regarding pCC3, the overexpression of collagen VI members was observed, including many collagen VI interacting proteins^35^. In general, the closer the relationship between survival outcome and pathways was, the greater the number of abundance changes in the pathway proteins. For instance, the increased immune tolerance in pCC1 could result in higher survival rates, while the upregulated collagen VI proteins in pCC3 worsened metastasis statuses due to the decellularization of extracellular matrix proteins. Thus, the proteogenomic evidence indicated another new classifier of the molecular subtypes of ESCC.

**Figure 5.**
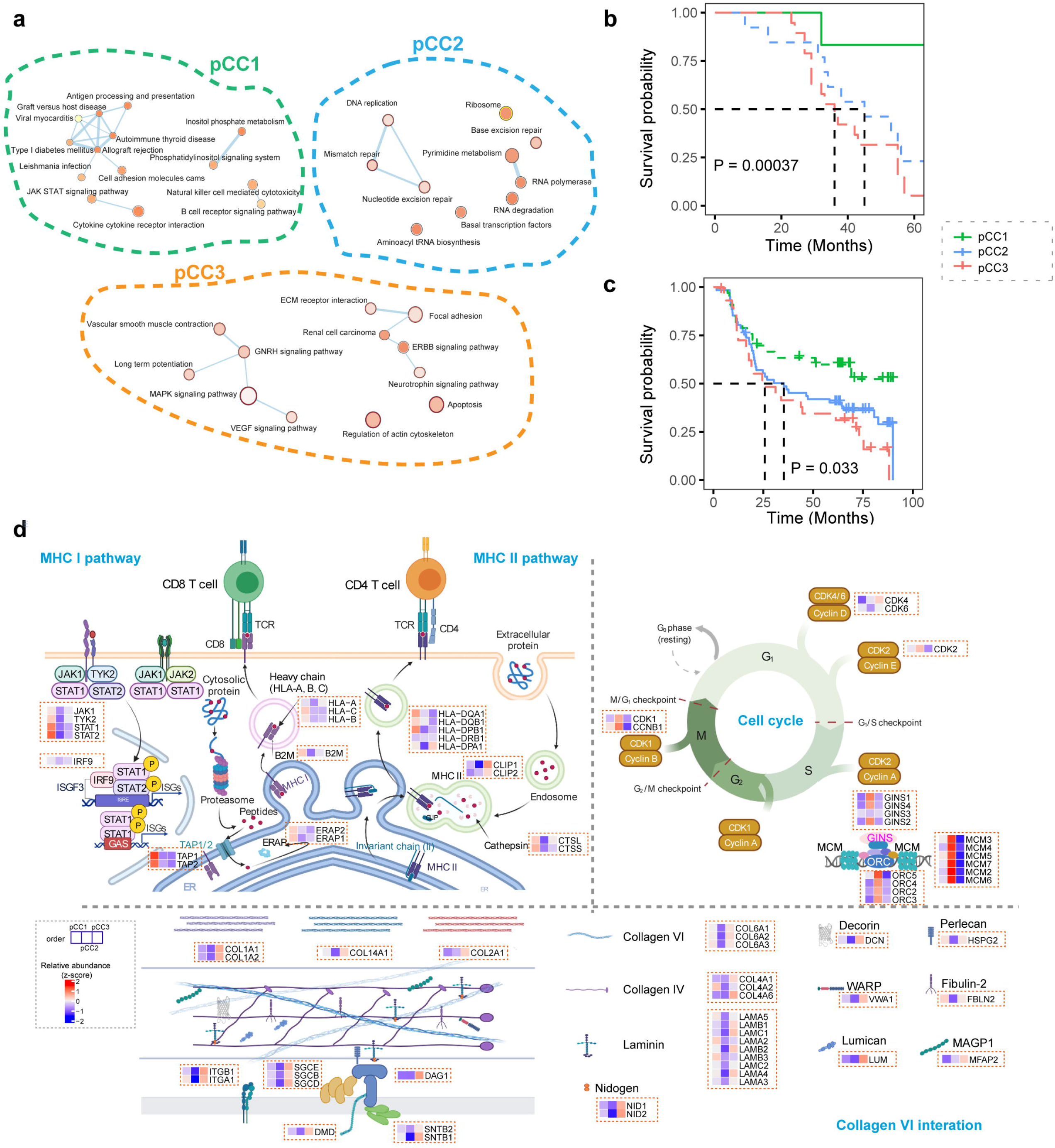
Proteomic subtypes of ESCC and subtype-specific pathway enrichment and represented signatures. (**a**) Enrichment maps of pathways and processes were determined by an integrative analysis of protein abundance in different ESCC molecular subtypes. (**b**) Kaplan–Meier curves of overall survival (OS) for each proteomic subtype of the ESCC cohort. P values were calculated by a two-sided log-rank test. (**c**) Kaplan–Meier curves of overall survival (OS) for each proteomic subtype of the public ESCC cohort. P values were calculated by a two-sided log-rank test. (**d**) Pathways activated in proteomic molecular subtypes of ESCC.

### Potential therapeutic strategies for the molecular subtypes of ESCC

There are three main strategies for ESCC therapy in the clinic—radiotherapy, chemotherapy and targeted therapy **(Fig.6a)**, even though some drugs in these therapy categories are not used for ESCC^36^.

**Figure 6.**
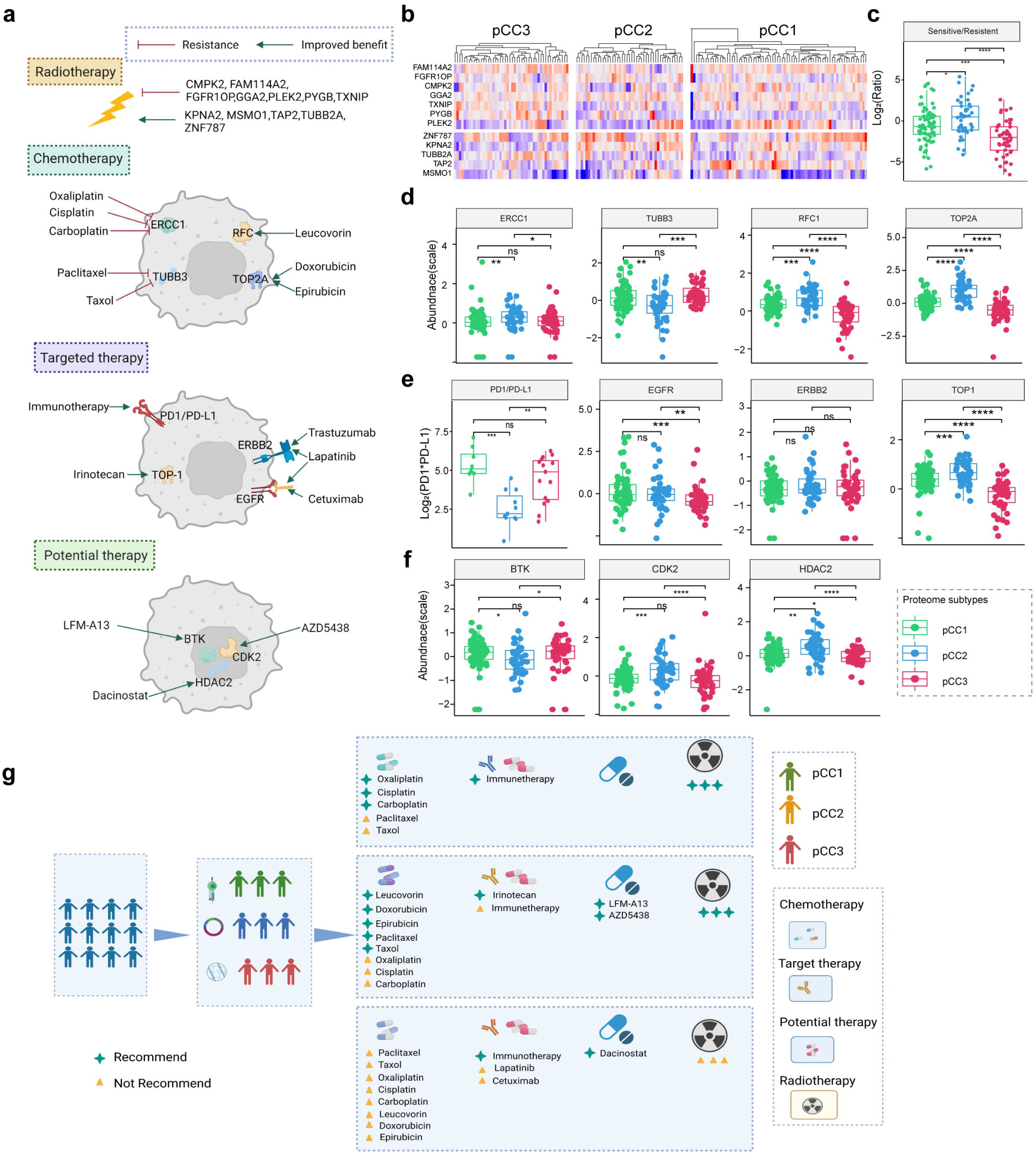
Therapeutic analysis for different molecular subtypes of ESCC. (**a**) Proteins related to different therapeutic strategies for ESCC. (**b**) Heatmap based on abundance distribution for radiosensitivity and resistance genes in different molecular subtypes. (**c**) Boxplot of the abundance ratio of radiosensitive vs. resistance genes in different molecular subtypes. (**d**) Boxplot of abundance distribution for drug targets related to chemotherapy in different molecular subtypes. (**e**) Boxplot of abundance distribution for target therapy in different molecular subtypes. (**f**) Boxplot of abundance distribution for potential targeted therapy in different molecular subtypes. (**g**) Summary of recommended therapeutic strategies for four molecular subtypes of ESCC.

Regarding radiotherapy for ESCC, alterations in protein abundance in ESCC tissues are positively correlated with radiotherapy sensitivity, such as changes in KPNA2, MSMO1, TAP2, TUBB2 and ZNF787 levels, whereas changes in the abundance of other proteins may strengthen resistance to radiotherapy, such as changes in FAM114A2, PYGB, SARS2, TXNIP, ATP4A, CMPK2, GGA2, PLEK2 and FGFR1OP levels^37^. A total of 12 proteins typically related to radiotherapy were identified in this study, and their abundance distribution in the ESCC subtypes derived from the DEPs mentioned above was hierarchized as shown in **Fig.6b**. We used radiotherapy-related response ratio by division of the abundance of radiosensitive proteins by that of the 5 radio-resistant proteins for evaluation, the higher values indicating the greater sensitivity to radiotherapy. As shown in **Fig.6c**, since the radiotherapy-related ratio of pCC3 was significantly distinct from that of PCC1 or pCC2, suggesting that the ESCC patients in pCC3 might be highly resistant to radiotherapy, whereas the patients in pCC1 or pCC2 might have better outcomes after radiotherapy.

Regarding chemotherapy in ESCC, the abundance of some proteins has been reported to be drug dependent^38^. ERCC1 is associated with resistance to oxaliplatin^39^, cisplatin^40^ and carboplatin^41^, TUBB3 is resistance to paclitaxel and Taxol^42^, and RFC^38^ and TOP2A^43^ are related to the effectiveness of leucovorin, doxorubicin and epirubicin. The statistical assessment of the abundance of the 4 drug-dependent proteins was conducted in the molecular subtypes of ESCC (**Fig.6d)**. Compared with the patients classified as pCC1 and pCC3, the patients classified as pCC2 exhibited higher ERCC1 and lower TUBB3 abundance, suggesting that they would be resistant to oxaliplatin, cisplatin and carboplatin and sensitive to paclitaxel and Taxol. The patients classified as pCC3 shared a lower abundance of RFC and TOP2A, significantly different from those classified as pCC1 and pCC2, implying that they might be resistant to leucovorin, doxorubicin and epirubicin (**Fig.6d**).

Regarding targeted therapy for ESCC (**Fig.6e**), the target genes are broadly divided into two types: traditional genes, including immunotherapy targeting PD1/PD-L1, trastuzumab and lapatinib targeting ERBB2, irinotecan targeting TOP-1, and lapatinib and cetuximab targeting EGFR, and potential genes, including LFM-A13 targeting BTK, AZD5438 targeting CDK2 and dacinostat targeting HDAC2^44^. All statistical comparisons of the target abundance in the three ESCC molecular subtypes are presented in **Fig.6e** and **6f**. The abundance of PD1 and PD-L1 was lower in pCC2 than in pCC1 and pCC3, indicating less immunotherapy benefit among these patients. The abundance of EGFR and TOP1 in pCC3 was significantly lower than that in other subtypes, implying that pCC3 patients might be less responsive to irinotecan, lapatinib and cetuximab. Regarding the potential targets, pCC2 patients had a higher abundance of CDK2 and HDAC2 than those classified as pCC1 and pCC3, which tended to be sensitive to AZD5438 and dacinostat, while they had a lower abundance of BTK, leading to less sensitivity to LFM-A13 (**Fig.6f**). The appropriate therapeutics for the three molecular subtypes of ESCC are summarized in **Fig.6g**.

### ESCC-related neopeptides discovered by the proteogenomic approach

Gene expression analysis of ESCC, either transcriptome or proteome, can reveal new peptides that are resulted from alternative splicing or mutated transcripts driven by cancer. Based on the integration of transcriptomic and proteomic analysis with a strict criterion for MS/MS signals, 10 top ESCC-related peptide candidates were selected for further evaluation (**Supplementary Fig.7**). The typical MS/MS spectra of neopeptides with convincing evidence of peptide fragmentation, AS2 and NT6, are presented in **Fig.7a**. The advanced assessment was based on three considerations: identification frequency in ESCC tissues, MHC affinity of these neopeptides and peptide abundance in molecular subtypes.

**Figure 7.**
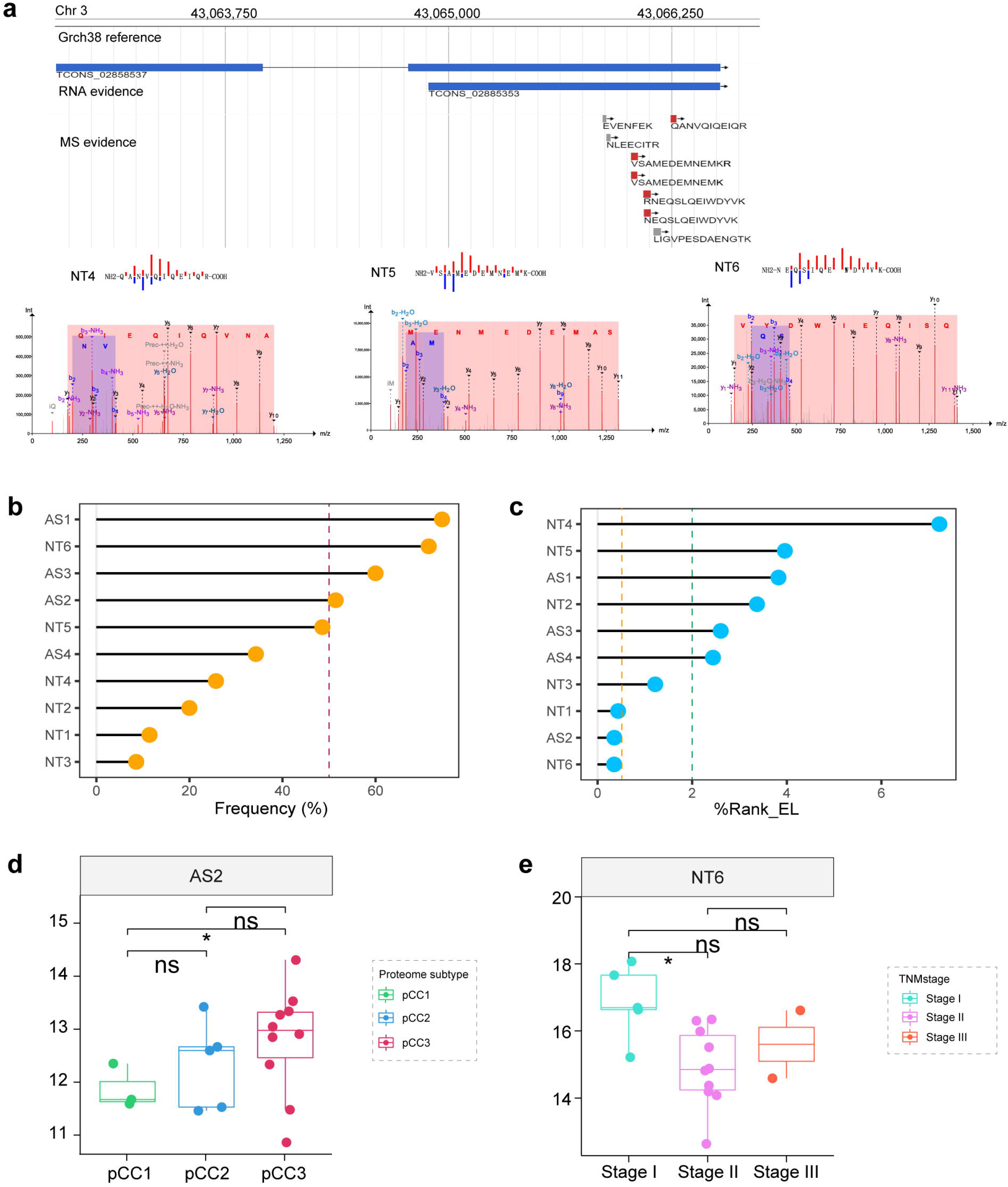
Novel peptides of ESCC. (**a**) Representative MSMS for AS2 and NT6. (**b**) Distribution of identified percentages of new peptides in ESCC tumor samples. The red dashed line represents the threshold at 50%. (**c**) Distribution of identified percentages for new peptides in ESCC tumor samples. The red threshold is 50%. (**d**) Boxplot of the abundance of AS2 across different molecular subtypes. (e) Boxplot of the abundance of NT6 across different TNM stages.

Among 46 paired ESCC tissues, all 10 neopeptides were identified in the tumor tissues but not in the paired adjacent tissues. The detection frequencies of the peptides are summarized in **Fig.7b**, indicating 4 neopeptides with over 50% detection rates; 74.3% in AS1, 51.4% in AS2, 60.0% in AS3 and 71.4% in NT6. A potential MHC affinity to peptides was analyzed by NetMHCpan 4.1^45^, a predictor based upon classified HLA alleles with the criteria of Rank_EL values less than 0.5 indicating stronger binding or 0.5 to 2 indicating weak binding. As shown in **Fig.7c**, NetMHCpan 4.1 predicted 3 neopeptides with strong binding affinity to MHC, AS2, NT1 and NT6, and only NT3 was a weak binder. With filtration of the detection frequency and MHC affinity, two neopeptides, AS2 and NT6, were identified as potential ESCC neoantigens. The abundance of AS2 and NT6 in different molecular subtypes was further assessed. The AS2 abundance in pCC3 was obviously higher than that in pCC1, and there was no difference in the NT6 abundance among the subtypes. The abundance of NT6 was higher in TNM stage I than in stage III. As pCC3 patients showed low sensitivity to radiotherapy and small molecular drug targeted therapy, AS2 is a potential immune therapeutic neoantigen for treating pCC3 patients.

## Discussion

We conducted a proteogenomic study of ESCC that concentrated on two sets of omics data, ESCC-related transcriptomes and proteomes. It is accepted that cells and tissues demonstrate a relatively poor correlation between the abundance of mRNAs and proteins, with an approximate correlation coefficient of 0.3-0.45. Why did we choose these two types of omics data for our proteogenomic study? First, gene expression status, either transcription or translation, is perceived to be strongly associated with cancer generation and development. Using information from two sides provides a comprehensive picture of cancer-related gene expression. Second, the comparison of gene expression status between tumor and adjacent tissues is generally conducted by analyzing DEGs and DEPs or using WGCNA^17^. These two approaches evaluate omics data from different angles, converging at significantly different abundances of gene expression or weighing the coexpressed genes. We attempted to combine the two approaches and deepen the integration of cancer-related differential gene expression data and WGCNA^17^ modules by studying mRNAs and proteins. As illustrated in Figures 3 to 5, the ESCC signatures were derived from the ESCC-related modules and hub genes, while ESCC molecular subtypes could be categorized based on DEGs and DEPs. Third, the informatic analysis of proteogenomic data should not be limited to the experimental data generated in the relevant studies; instead, it should be extended to the available data in the published datasets, promoting the integration of multiple omics datasets and verification of the evidence by others. Evaluating the TCGA dataset and the modules derived from ESCC proteogenomic data revealed ME5 as the module specific for both ESCC and EAC and ME15 as the only module specific for ESCC alone. ESCC molecular subtypes were not only inferred from proteogenomic data but also confirmed by other independent ESCC proteome datasets. Combining transcriptomic and proteomic datasets from different cohorts and multiple informatics datasets provided novel insight into ESCC biology and potential new options for clinical application.

Exploration of the consensus module analysis of ESCC-related proteogenomic data revealed that EGME5 is an ESCC-specific module in which esophageal tissue-specific genes are enriched. Furthermore, the combinations of the 8 genes could enable the discrimination of ESCC from other cancer types at either the transcript (Figure 3d) or protein level (**Fig.3e**). In this study, WGCNA^17^ demonstrated how powerful the approach to the ESCC-related gene network was and how useful the panel of esophagus-specific proteins could be in clinical applications. Immunohistochemical analysis of SPRR3 revealed that it was significantly downregulated in ESCC and associated with differentiation grade^46^. S100A14 is one member of the S100 protein family that comprises 25 members; these two represent widely accepted tumor biomarkers. The abundance of S100A14 in ESCC tissues is downregulated compared with their corresponding adjacent normal tissues, and lower S100A14 expression predicts poorer overall survival^47^. Thus, other observations regarding ESCC-dependent gene expression supported the ESCC-related modules reasonably deduced from proteogenomic data.

Cancer characteristics are heterogeneous; therefore, finding an appropriate classification method for cancer patients, especially from a molecular perspective, is urgently required in oncological clinics and research. Several studies on the molecular characterization of ESCC have been documented. Lin et al. performed integrative clustering of DNA methylation, mRNA, somatic copy number and microRNA to classify ESCC into two subtypes, ESCC1 with alterations in the NRF2 pathway and ESCC2 with higher rates of mutation of NOTCH1 or ZNF750 and greater leukocyte infiltration^48^. Based on transcriptome data, Liu et al revealed three ESCC subtypes associated with a distinct prognosis and lymph node metastasis in ESCC patients as the subtypes of metabolic, inflammatory and metastatic, and cell proliferation^49^. By combining ATAC-seq data with the RNA-seq data of ESCC, Li et al established a survival-related subtype classifier called PrSC, which captured stromal heterogeneity in ESCC and predicted the clinical response to immunotherapy^50^. There are limited reports regarding ESCC subtypes derived from proteomic observations. With label-free phosphoproteomics, Li et al employed consensus clustering for the top 25% of the most variable proteins to stratify two ESCC subtypes, and the patients in S2 had a worse 5-year overall survival rate and disease-free survival outcomes than those in S1^51^. The classification processes for ESCC mentioned above are unsatisfactory due to their poor clinical significance; thus, other methods involving detection in plasma or that provide guidance for therapy are required. In contrast to other reports, we proposed a new ESCC category, pCC1, as an immune-related subtype with activated M1 macrophages, pCC2, as a cell cycle-related subtype, and pCC3, as a metastasis-related subtype. These new types of ESCC classification exhibit their own features: 1) the molecular subtypes were not only derived from the DEP clustering in this study but were also supported by the other public dataset of ESCC proteomes; 2) such proteome-based classification system subtyped ESCC based on the transcriptome; thus, transcriptional signals could partially represent the subtyping ESCC; 3) the ESCC-related DEPs for ESCC subtypes largely overlapped with the ESCC-dependent hub genes identified by WGCNA, specifically those in ME5, while some key proteins encoded by these hub genes were detectable in plasma, implicating the plasma biomarkers of ESCC as a reference to determine patient subtypes; and 4) the different molecular subtypes of ESCC favored the selection of therapeutic recommendations. The proteome-based view affords a new and clinic-feasible categorization process for ESCC.

## Methods

### Specimens and clinical data

All samples for the current study were obtained through the Biobank of Anyang Cancer Hospital. All cases with pathologic diagnoses for tumor-node-metastasis (TNM) stages were evaluated on the basis of the Cancer Stage Manual, 7th ed., issued in 2009 by the American Joint Committee on Cancer^52^. All specimens that matched adjacent normal esophageal tissues were used as controls. After surgery, the fresh and paired tissues, tumor and adjacent, were washed with cool phosphate buffer saline, immediately snap frozen in liquid nitrogen, and stored at −80 °C until transcriptomics and proteomics analysis. A written, informed consent was signed by patients, and the procedure of sample collection was approved by the ethics committee of Anyang Cancer Hospital.

### Sample processing for mRNA sequencing

The frozen tissues were homogenized and divided into two aliquots for RNA and protein extraction. Samples that passed RNA quality control and had a minimum RIN (RNA integrity number) score of 7 was subjected to RNA sequencing. The raw Illumina sequence data were demultiplexed and converted to FASTQ files, and samples were then assessed for quality by mapping reads to the hg38 human genome reference.

### Protein extraction and tryptic digestion

The frozen tissues were ground with liquid nitrogen, and the powdered tissues were homogenized by sonication in lysis buffer containing 2% SDS, 7 M urea, 10 mM EDTA, 10 mM PMSF, 10 mM DTT, and 0.1 M Tris-HCl, pH 7.6. The homogenized tissues were centrifuged at 20000g, and the resulting supernatants were reduced by 10 mM DTT and alkylated by 55 mM iodoacetamide. Total protein content was measured using the Bradford assay. The treated proteins were tryptic-digested following the FASP protocol as described by Mann ^53^.

### LC-MS/MS analysis

Each sample was resuspended in buffer A (5% ACN and 0.1% FA in water) and centrifuged at 20,000 g for 10 min. For samples quantification in data independent acquisition (DIA) mode, peptides separated from nanoHPLC by an analytical C18 column (75μm*35cm*1.7μm, in-house) and subjected into the tandem mass spectrometry Q EXACTIVE HF (Thermo Fisher Scientific, San Jose, CA) for detection. The gradient was run at 500 nL/min starting from 5 to 25% of buffer B (95%ACN, 0.1%FA) in 150 minutes, going up to 35% in 10 minutes, then going up to 80% in 5min, then maintenance at 80% B for 5 minutes, and finally return to 5% in 0.1 min and equilibrated for 7 min. The parameters for MS analysis are listed as following: electrospray voltage: 1.6 kV; precursor scan range: 400-1250 m/z at a resolution of 120,000 in Orbitrap; MS/MS fragment scan range: >100 m/z at a resolution of 30,000 in HCD mode; The DIA method was set with 45 windows from 400-1250 m/z and the isolation window was set 17m/z. The stepped collision energy setting: 22.5, 25, 27.5; Automatic gain control (AGC) for full MS target and MS2 target: 3e6 and 1e6, respectively.

### Quantification of DIA proteomics data

For DDA data, all raw files were combined and searched by MaxQuant ^54^ using the swissprot database of human as reference. The identification files were imported into Spectronaut (version 12) software as DIA library, and all DIA data were processed with this library. At the proteome level, the ion library was generated from the pooled ESCC and adjacent samples in DDA mode, including 149,527 peptides corresponding to 11,548 protein groups. In the 46 paired ESCC tissues, 105,499 peptides corresponding to 10,067 proteins were identified with DIA mode (detailed identified protein number for each sample depicted in Figure S1a); 7,985 proteins were present in at least 20% of the samples (i.e., quantifiable proteins).

### Parallel reaction monitoring analysis

New peptides detected in samples by the DIA analysis for further validation by targeted proteomic analyses using parallel reaction monitoring (PRM) and processed the targeted method by skyline^55^ in the pooled samples of all ESCC patients. All peptide separations were performed using the same LC gradient as mentioned in the DIA setting. PRM analyses were performed on a Q-Exactive HF mass spectrometer (Thermo Fisher Scientific). The SIM scan event was collected using a m/z 380–1,500 mass selection, an Orbitrap resolution of 17,500 (at m/z 200), target automatic gain control (AGC) value of 3 3 106 and a maximum injection time of 30 ms. The PRM scan events used an Orbitrap resolution of 17,500, an AGC value of 1 3 106 and maximum fill time of 80 ms with an isolation width of 2 m/z. Fragmentation was performed with a normalized collision energy of 27 and MS/MS scan were acquired with a starting mass of m/z 150. All PRM data analysis was performed using Skyline^55^ software. Validation was achieved by comparing the fragment ion ratios and retention times of the endogenous variant peptide to that of the synthesized peptide standards.

### WGCNA analysis of transcriptome and proteome datasets

We performed further quality-control on the matrix of normalized expression values to remove any transcripts with either zero variance or a missing value prior to performing weighted gene coexpression network analysis (WGCNA)^16^. We implemented the WGCNA package to create a weighted adjacency matrix. This weighted adjacency matrix was used to generate a topological overlap matrix (TOM) and dendrogram. A dynamic hybrid branch cutting method was implemented on the resulting TOM-based dendrogram to identify modules. A cut height of 0.3 was set to merge module eigengenes (ME; first principal component of each network) that have a correlation of 0.7 or greater. We then estimated MEs, which are the first principal components for each gene expression module after a singular value decomposition is performed on the TOM. To explore the consistent modules between ESCC and pan-cancer types, module preservation and quality statistics were computed with the modulePreservation function (nPermutations = 200) implemented in the WGCNA package^16^.

### Differential transcriptomic and proteomic analysis

Transcriptomic and proteomic data were used to perform differential analysis between groups of samples. A Wilcoxon rank-sum test was performed to determine differential abundance of mRNAs and proteins. At least four samples in both groups were required to have non-missing values and the p-value was adjusted using the Benjamini-Hochberg procedure. p-value for the mRNA or protein change was required to be less than 0.05.

### Gene set enrichment analysis

A Gene Set Enrichment Analysis (GSEA)^56^ was performed to identify pathways enriched in the Molecular Signatures Database (MSigDB)^57^ Hallmark gene set. A nominal p value of < 0.05 was considered statistically significant.

### Molecular subtyping based on transcriptome and proteome data

Consensus clustering was performed using the R package ConsensusClusterPlus^58^. Samples were clustered using Euclidean distance as the distance measure. The cumulative distribution function (CDF) of the consensus matrix for each k-value was measured and Clustering by k□ =□ 3 had the lowest proportion of ambiguous clustering (PAC). The relative change in area under the CDF curve increased 30% from 2 clusters to 3 clusters, while others had no appreciable increase. Taken together, proteome clusters were defined using k-means consensus clustering with k□ =□ 3. OmicsEV^15^ is an algorithm that estimates the significance of selected function terms based on gene expression. When functional prediction derived from mRNAs and proteins is close, the AUROC values for the two datasets remain in the diagonal region. Kaplan–Meier survival curves^59^ (log-rank test) were used for overall survival (OS) of the transcriptomic or proteomic subtypes and R package “survival” was used for survival statistical tests.

## Data availability

All data needed to evaluate the conclusions in the paper are present in the paper and/or the Supplementary Materials.

## Acknowledgements

We thank all the patients who participated. This work was supported by the National Program on Key Basic Research Project (2017YFC0906703).

## Author information

GH and SL contributed to the study concept and design. GH and SL were responsible for the data acquisition and analysis. SL and GH wrote the manuscript. FZ contributed to the collection of samples and clinical data, SX, ZR, ZJ, and YH contributed to critical revision of the manuscript. All authors revised and approved the final version.

## Supplementary information

Figure S1. Global overview of quantified mRNA and proteins among ESCC and adjacent tissues.

Figure S2. Function annotation of modules defined by WGCNA.

Figure S3. Cytoscape network for hub genes.

Figure S4. Representative molecular functions related with subtypes defined by mRNA expression.

Figure S5. Evaluation of molecular signatures related with clinical indicators at transcriptome level

Figure S6. Pipeline for potential neo peptide identification and verification

Figure S7. PRM validation of potential neoantigens of ESCC

Figure S8. Module abundance distribution for different ESCC molecular subtypes.

