## supplemental file for "Integration of proteogenomic analyses in esophageal squamous cell carcinoma"

Correspondence to:

**This PDF file includes: Supplementary Figure 1 to 8**

**SUPPLEMENTARY FIGURES**

**Supplementary Figure 1**

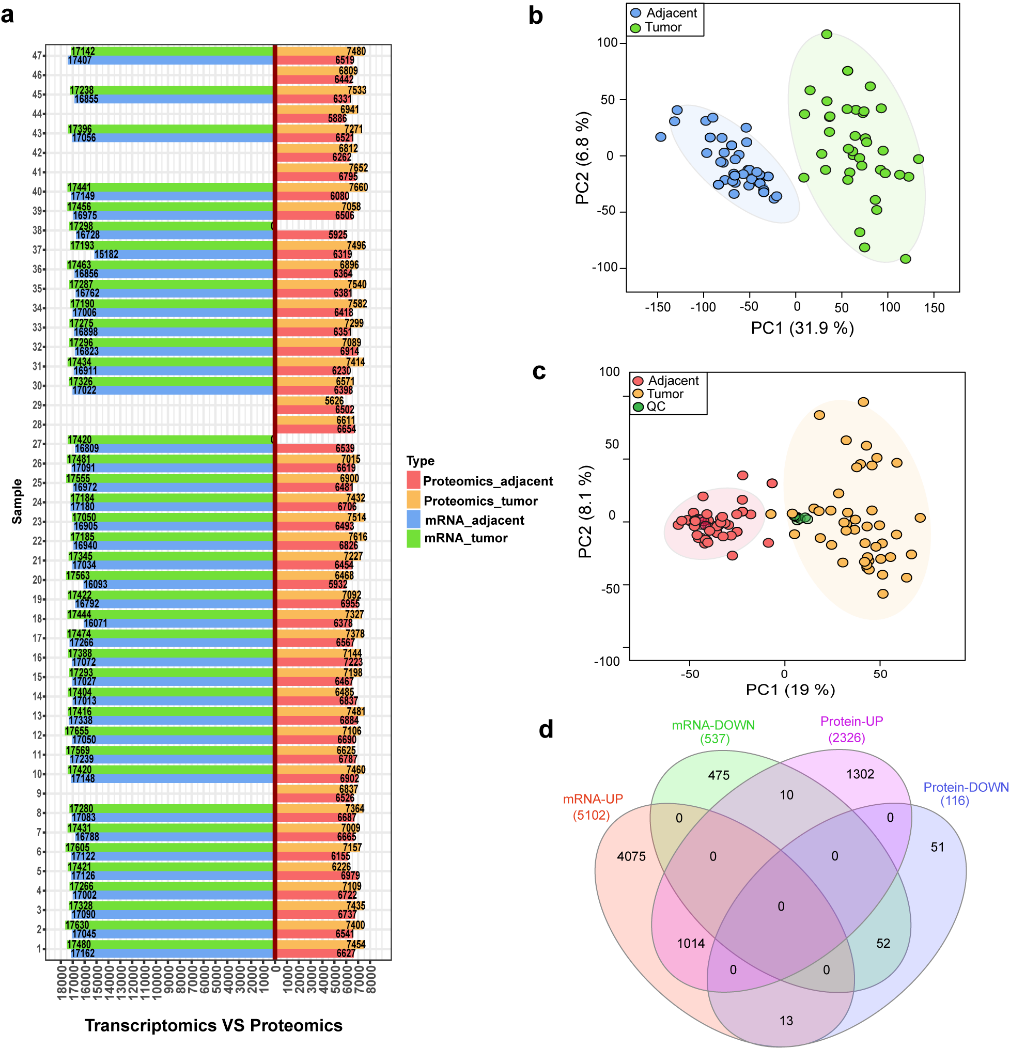

**Supplementary Figure 1. Global overview of quantified mRNA and proteins among ESCC and adjacent tissues.** (a) Identified mRNAs or proteins in each sample. (b) PCA distribution of ESCC and adjacent tissue samples using all transcriptome data. (c) PCA distribution of ESCC and adjacent tissue samples using all proteome data. (d) Venn plot based on mRNAs and proteins up regulated or down regulated in ESCC group compared with adjacent group.

**Supplementary Figure 2**

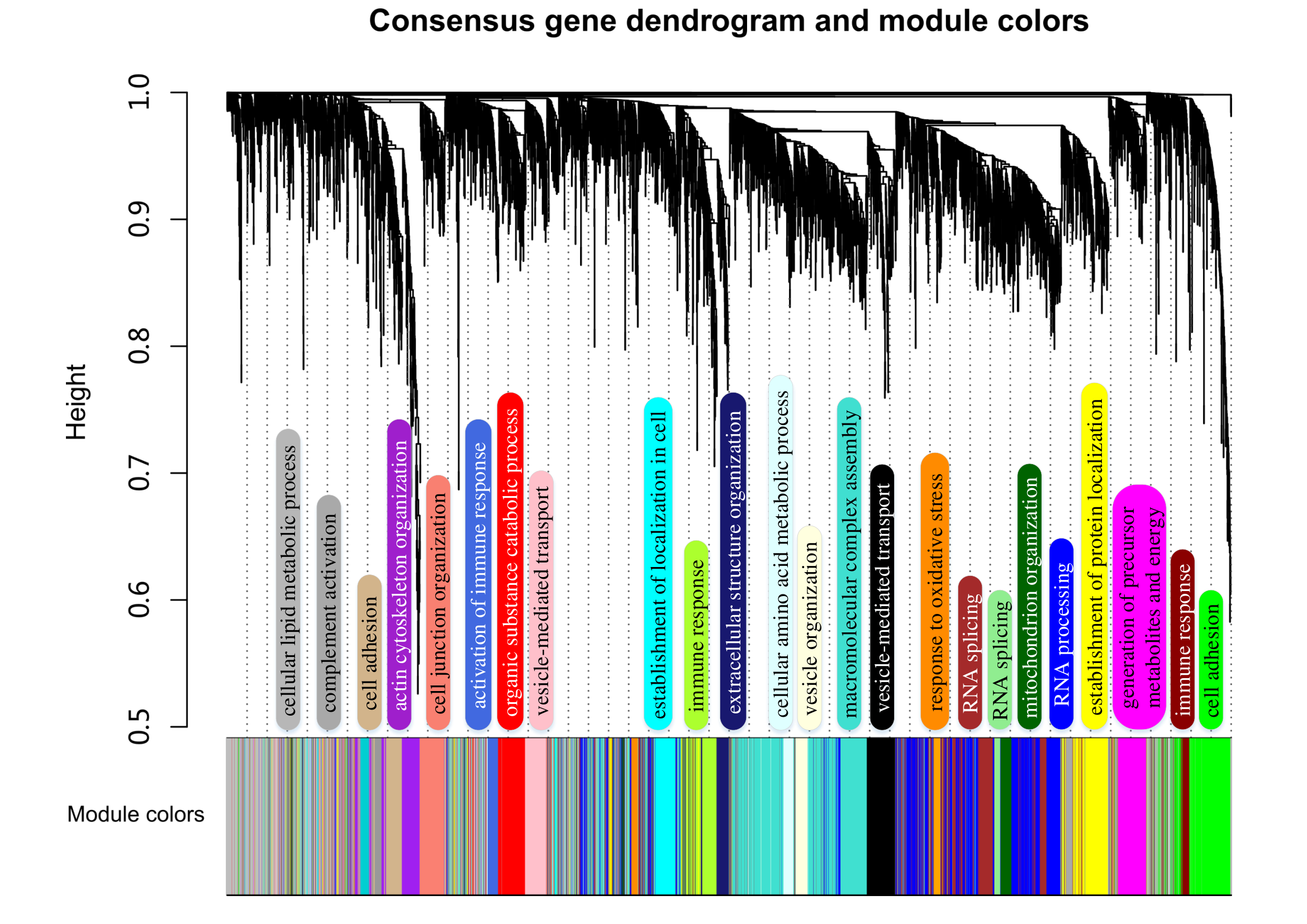

**Supplementary Figure 2. Function annotation of modules defined by WGCNA.** Cluster dendrogram and color representation of the coexpression network modules produced by average linkage hierarchical clustering of genes based on topological overlaps in the transcriptome and proteome data. Molecular function based for major modules were annotated.

**Supplementary Figure 3**

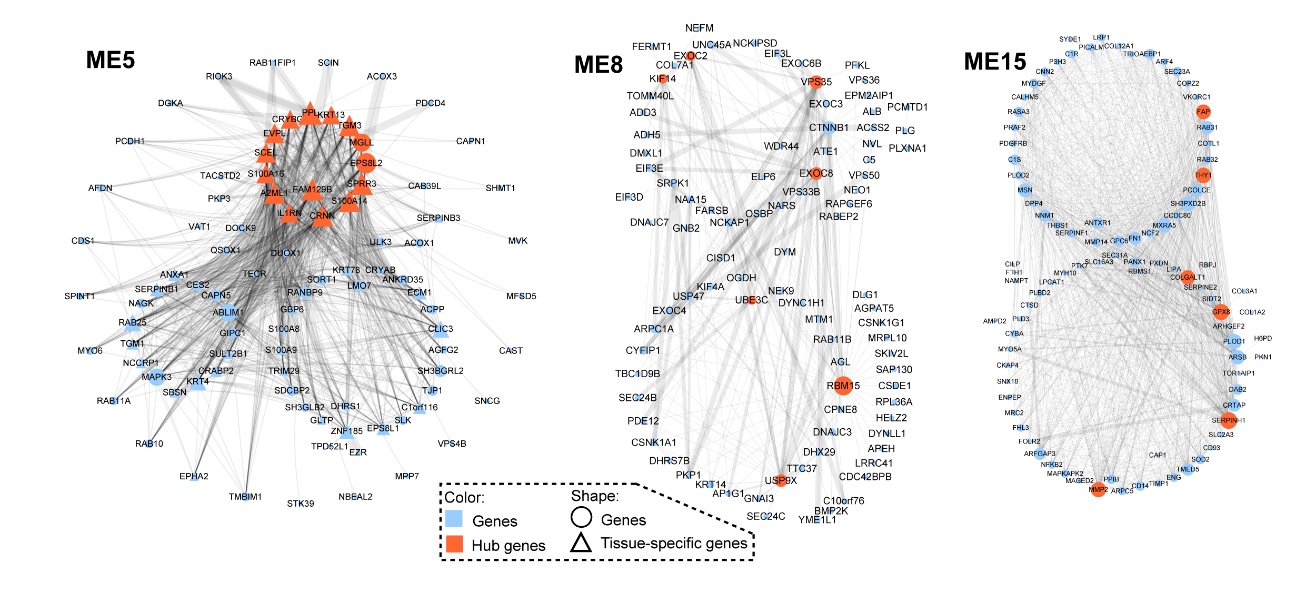

**Supplementary Figure 3. Cytoscape network for hub genes.** Hub genes were red colored and tissue-specific genes were represented with triangles.

**Supplementary Figure 4**

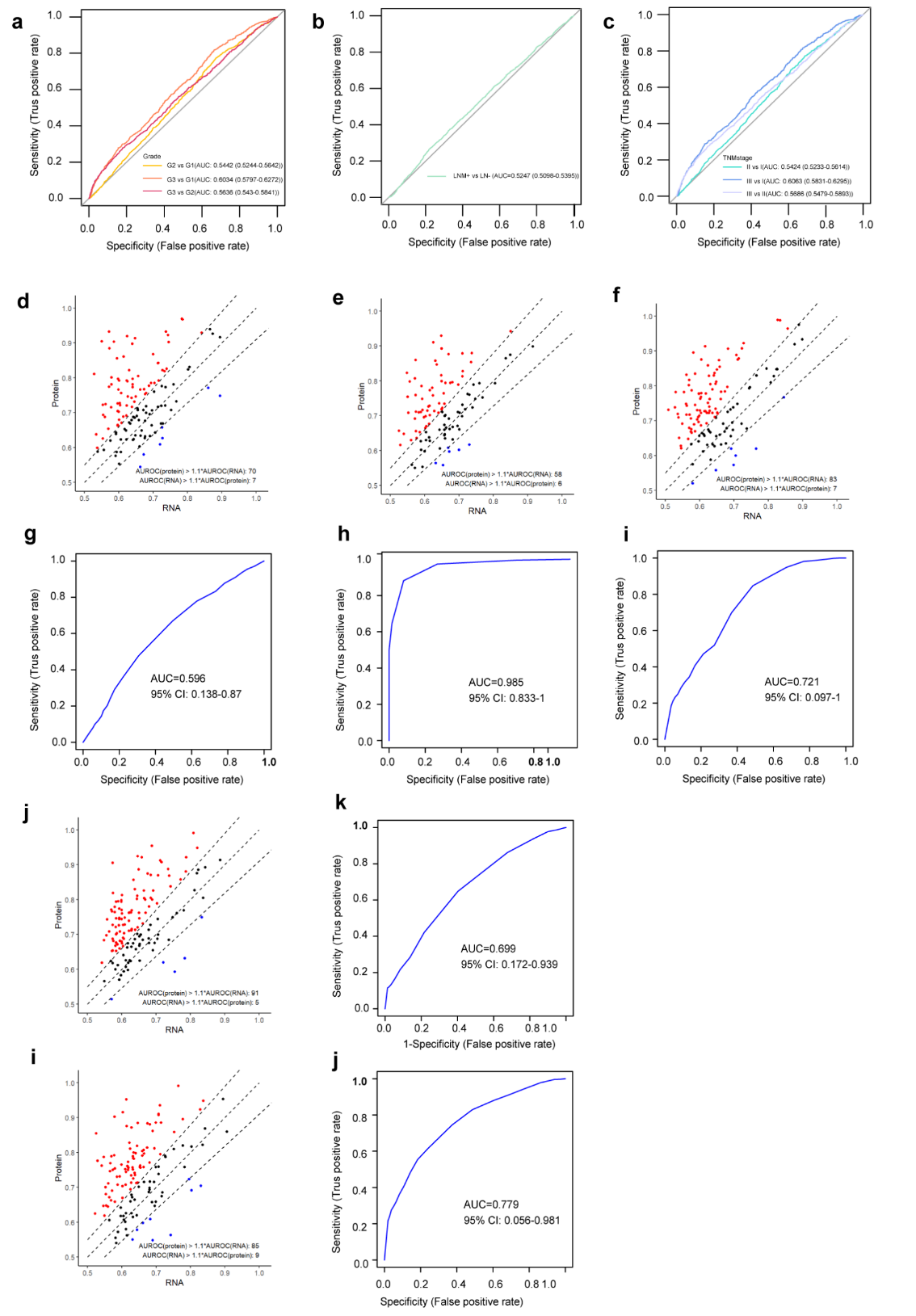

**Supplementary Figure 4. Representative molecular functions related with subtypes defined by mRNA expression.** Receiver operating characteristic (ROC) curves for the essential gene prediction classifiers on (a) grade, (b) LNM and (c) TNM stage datasets with combined mRNA features. Area under the receiver operating characteristic curve (AUROC) for KEGG (Kyoto Encyclopedia of Genes and Genomes) pathway membership prediction using RNA and protein data. Red and blue indicate pathways with >10% difference between the two. (d-j) represents prediction for Grades, LNM and TNM.

**Supplementary Figure 5**

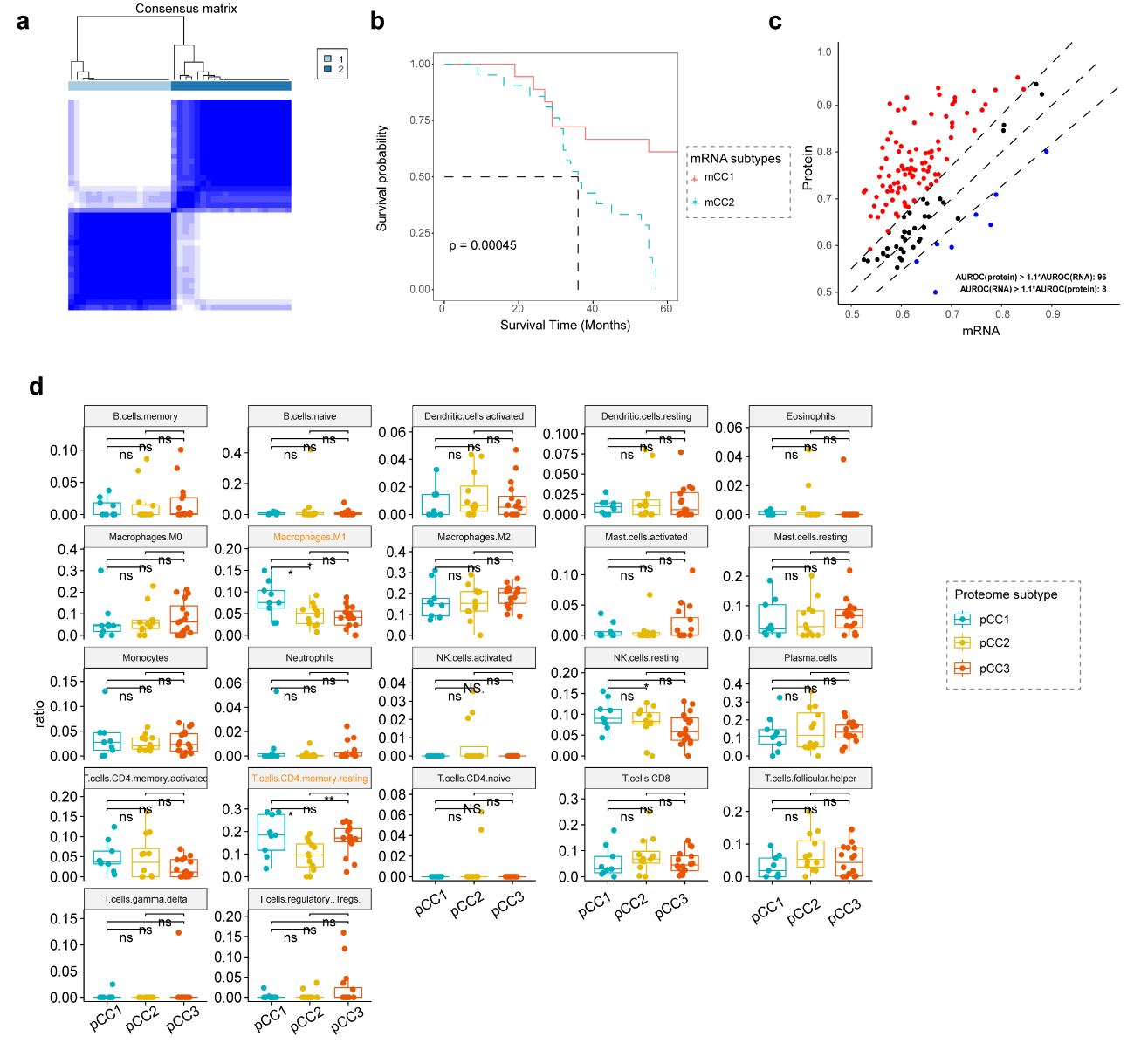

**Supplementary Figure 5. Evaluation of molecular signatures related with clinical indicators at transcriptome level.** (a) Consensus clustering matrix for k = 2. (b) Kaplan-Meier curve showed different prognosis between the two clusters. (c) Area under the receiver operating characteristic curve (AUROC) for KEGG (Kyoto Encyclopedia of Genes and Genomes) pathway membership prediction using RNA and protein data based on survival data. (d) Boxplot of single cell abundance based on proteome subtypes.

**Supplementary Figure 6**
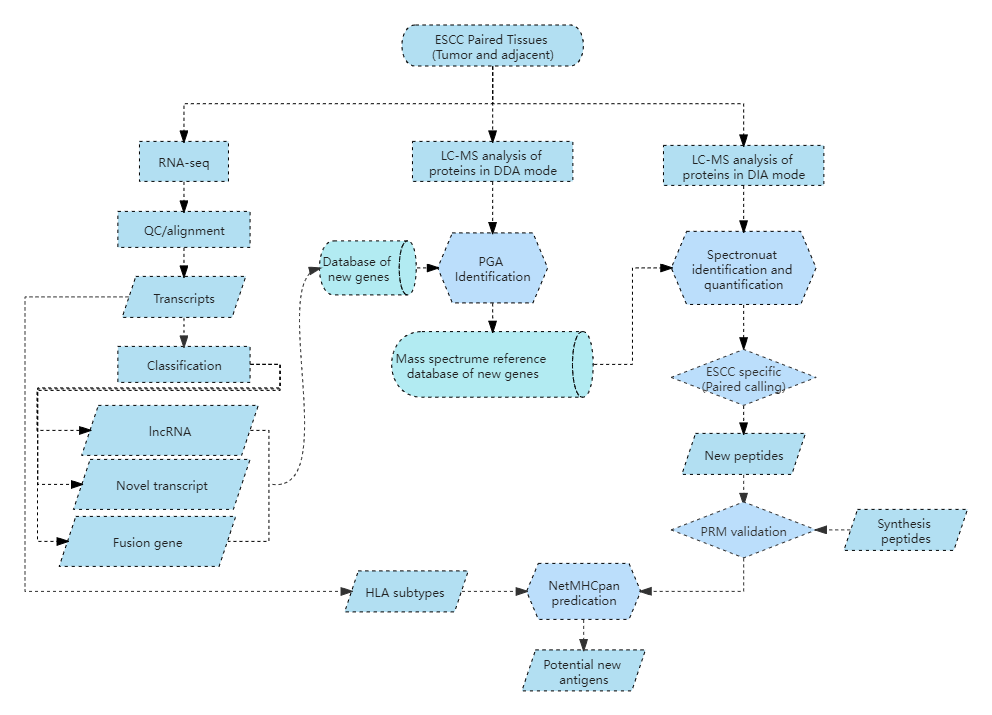

**Supplementary Figure 6. Pipeline for potential neo peptide identification and verification.**

**Supplementary Figure 7**

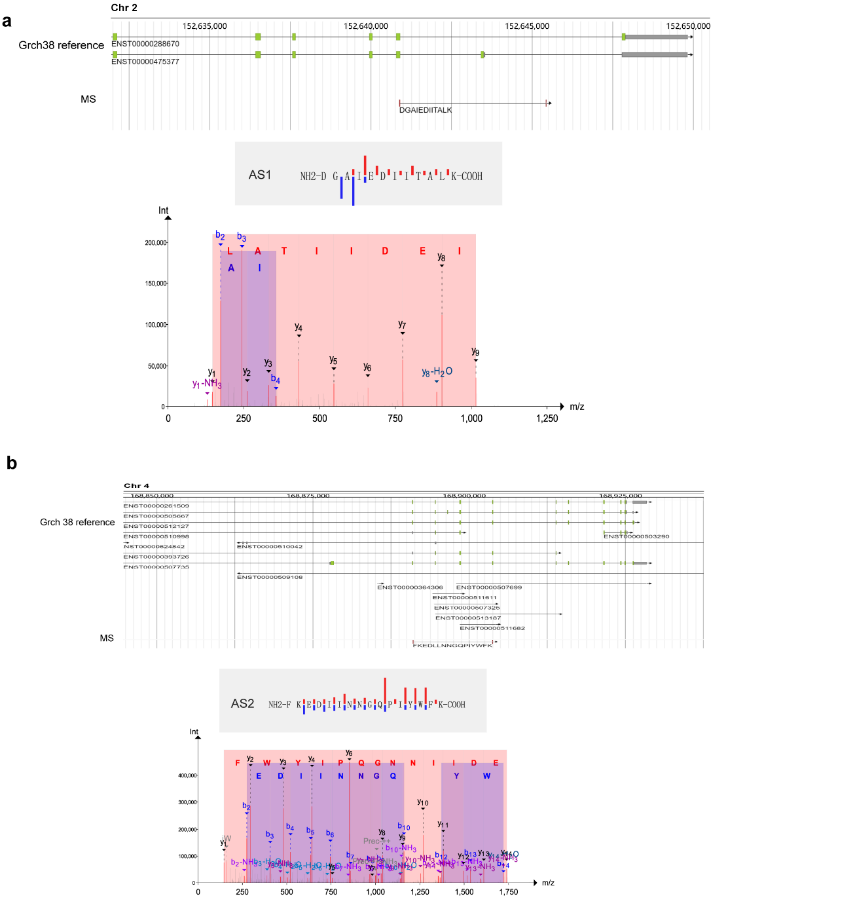

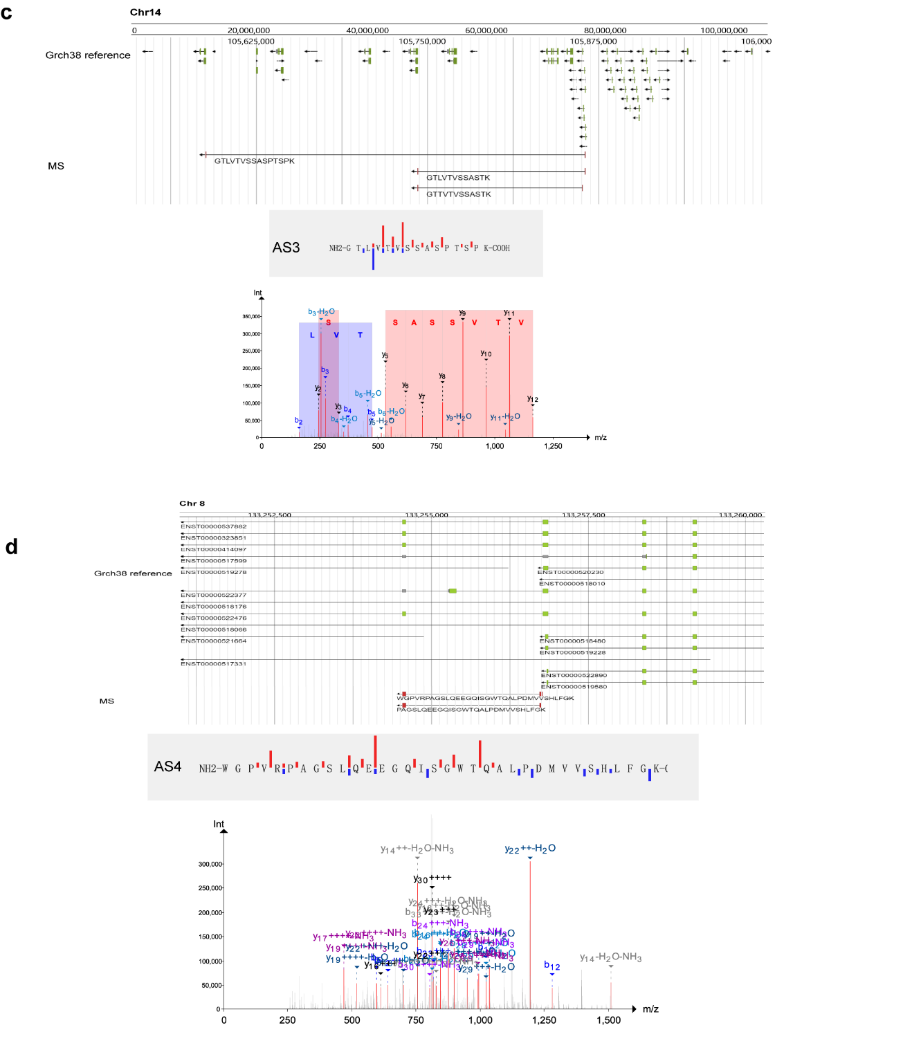

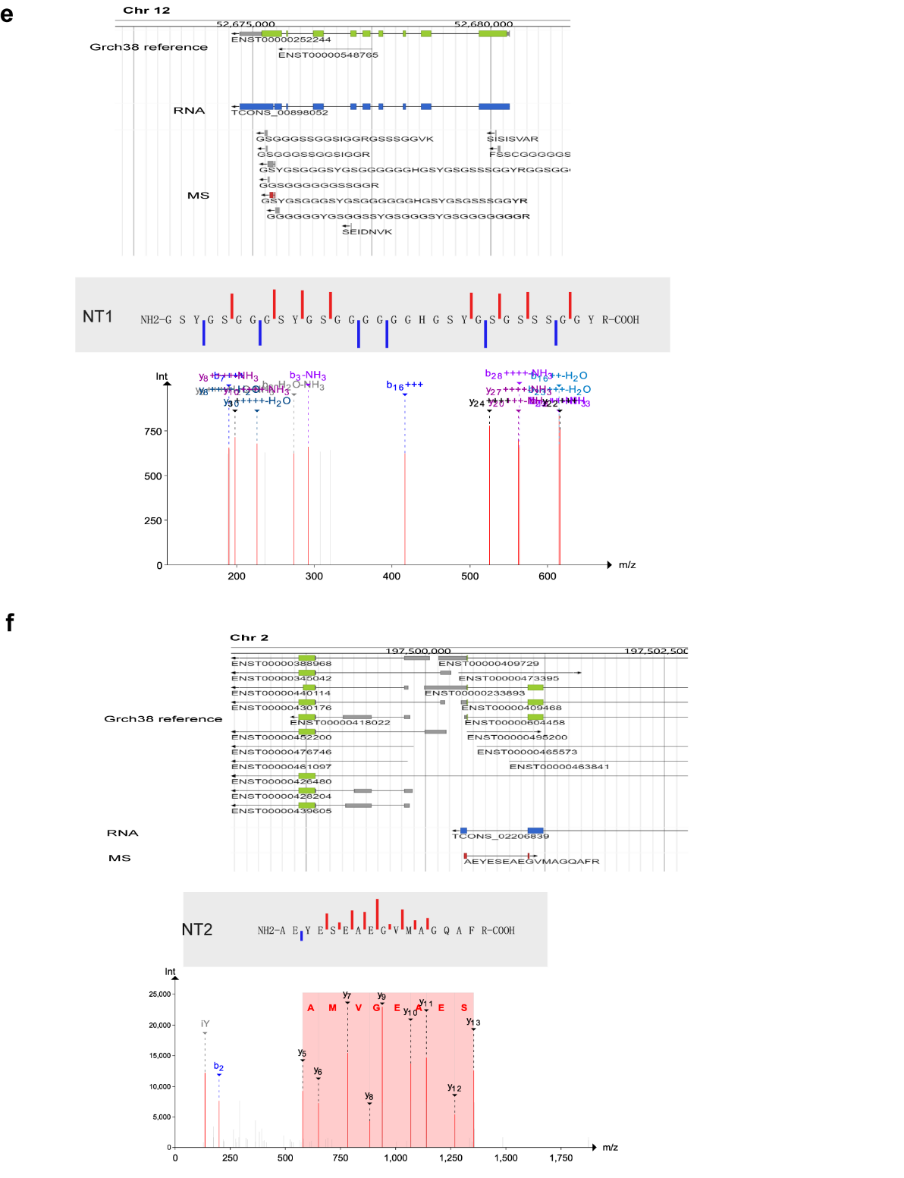

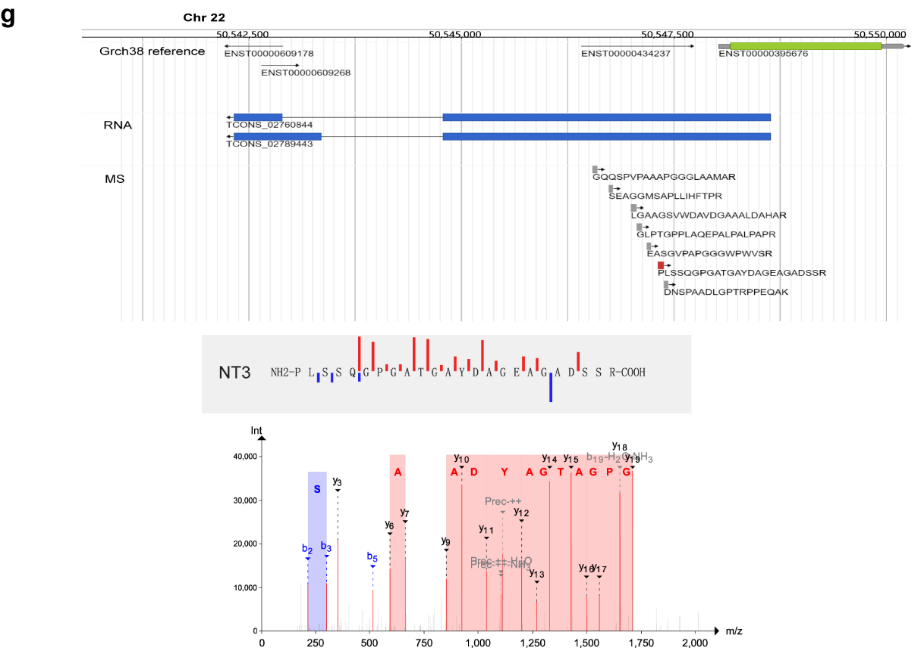

**Supplementary Figure 7. PRM validation of potential neoantigens of ESCC**

**Supplementary Figure 8**

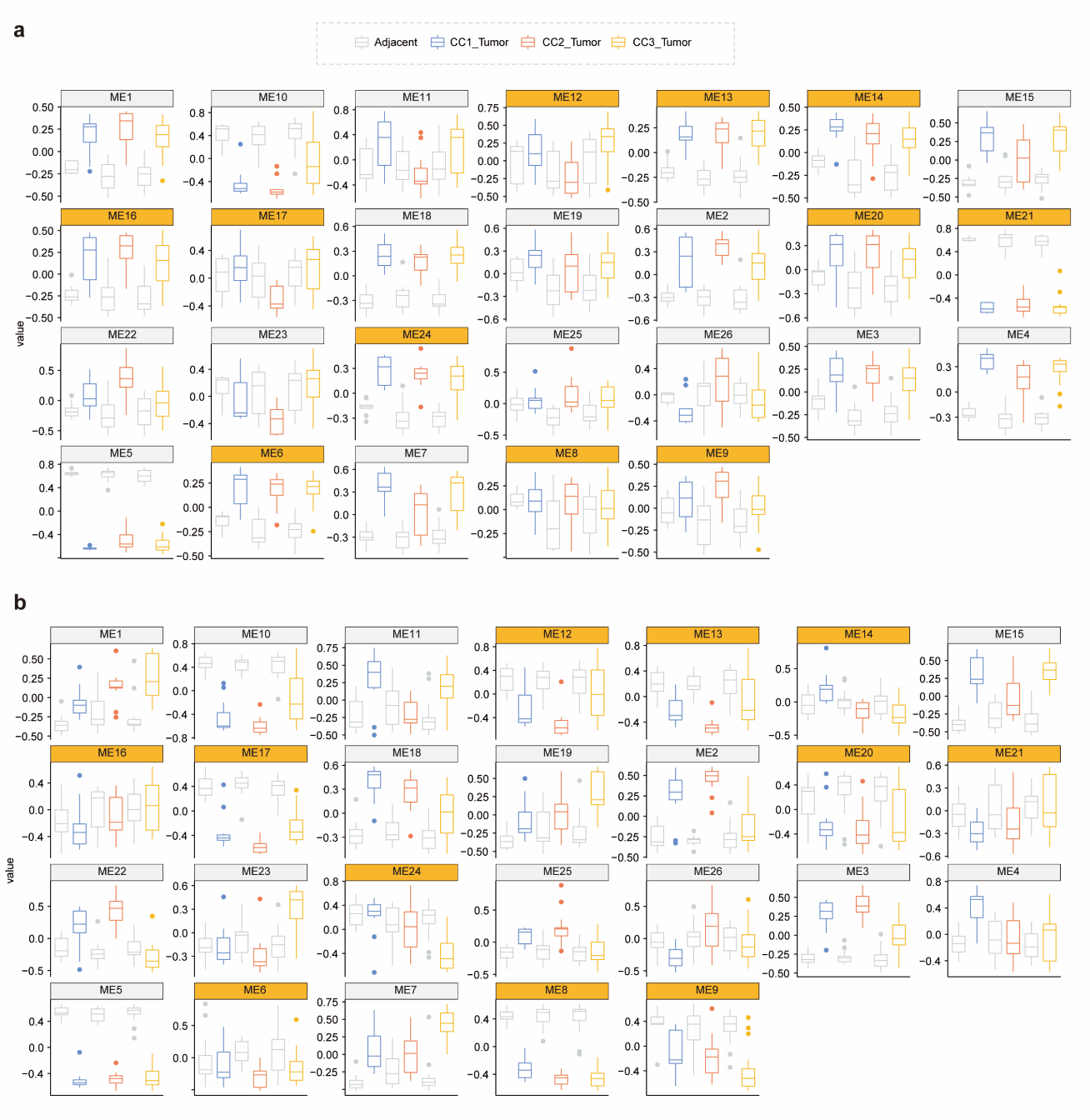

**Supplementary Figure 8. Module abundance distribution for different ESCC molecular subtypes.**
